# Remote and Independent Detection of Human Stress Using Sweat and AI

**DOI:** 10.64898/2026.01.21.700962

**Authors:** Anna Kochnev Goldstein, Yoav Goldstein, Yuri Feldman, Sharon Einav, Paul Ben Ishai

## Abstract

Previous studies exploring human sweat ducts as biological antennas in the sub-THz range have shown that the electromagnetic (EM) response of the skin is modulated by the person’s mental and physical stress. These findings naturally raised hopes of a new remote avenue for detecting human stress. However, as those studies unmasked stress using correlations with well-established markers such as the Galvanic Skin Response (GSR), the question of whether the EM response could serve as an independent marker of stress remained unanswered. Here, we provide a positive answer to this question by showing that machine learning models trained on EM reflections from 21 participants, subjected to physical and mental stress, were able to estimate the presence of stress in a signal from a new participant, in a matter of seconds, with above 90% accuracy.

## 1 Main

While stress itself is so deeply rooted in our biology and has such an intimate effect on our behavior, the tools available for detecting it and monitoring it are very limited. It is a sobering fact that in this modern age, we still cannot gauge the discomfort of an unconscious patient beyond a crude estimate of their pulse and blood pressure. A reliable and remote method for detecting and monitoring stress would be much welcomed.

When Optical Coherence Tomography (OCT) images revealed that the human sweat ducts are remarkably helical, a potential new candidate for such remote detection was born: The unexpected discovery led researchers to hypothesize that maybe if they bear such morphological resemblance to classical helical antennas, they might also function as such [1]. Subsequent OCT studies [2, 3] and computational models [4] showed that the center frequency for the axial mode for the human structures should lie in the sub-THz range of 360-520 GHz. As the human sweat ducts are also controlled by the sympathetic nervous system, which is responsible for our fight-or-flight response [5], this led to an intuitive subsequent hypothesis that the transmitted signals should carry information relating to human stress.

The latter hypothesis was addressed in several studies that showed that the EM emissivity and reflectivity of the human body in the sub-THz frequency range is modulated by the mental or physical stress experienced by the person under measurement [1, 6–17]. Naturally, many have wondered if these findings could be an avenue to remotely monitor personal stress. However, in those studies, the embedded information only became apparent through correlation with well accepted physiological parameters of stress, principally the Galvanic Skin Response (GSR). Obviously, if stress could be identified from the EM signal alone without correlations to other physiological measures, a new diagnostic avenue would be opened.

This work establishes this missing link. An EM signal in the range of 426-432 GHz was focused on the palms of 21 human participants and the reflection coefficient continuously recorded for 18 minutes while they interacted with a software application specifically designed to induce mental and physical stress. The recorded data was processed to create train and test features for supervised and unsupervised machine learning (ML) models with the goal to answer the following questions: (a) Can a model be trained to correctly distinguish between stress and no stress in a binary classification task? (b) Can a model distinguish between mental stress, physical stress, and no stress in a multi-class classification task?

## 2 Results

To answer the fundamental question of whether the EM recordings from the skin carry inherent information relating to the person’s stress, the employed supervised models (CNN and KNN with 3 neighbors), were tested on binary classification between “no stress” and “stress”, with no information on the type of stress (both mental and physical stress features were used but both types were given the same label). The accuracy of this binary classification as a function of signal length is depicted in Figure 1.

**Fig. 1:**
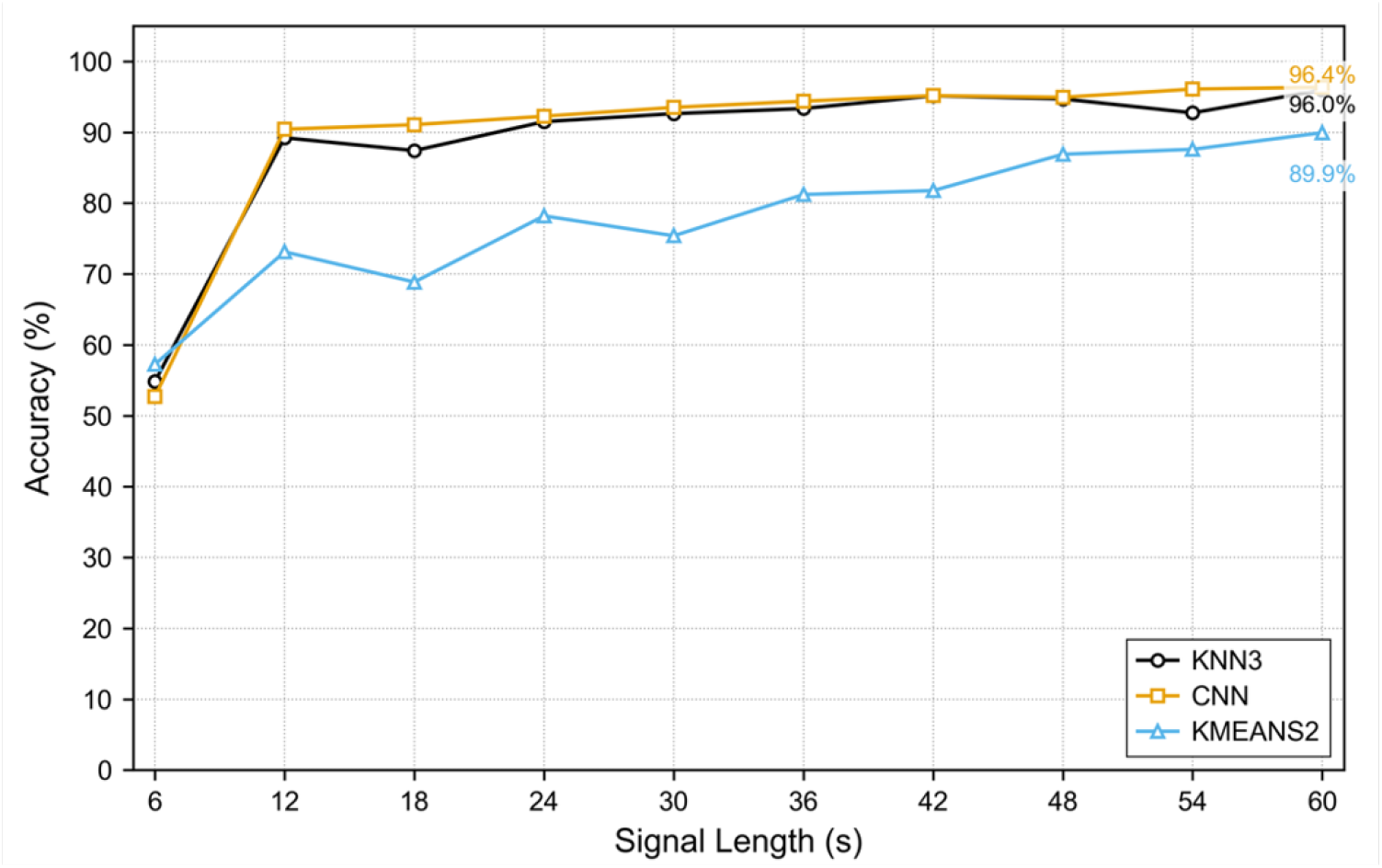
Binary classification accuracy of “stress” versus “no stress” features as a function of feature length (6-60 seconds) using the selected supervised models: KNN3 and a CNN, and the KMEANS2 unsupervised model.

Figure 1 shows that the supervised models were able to identify stress in a recording from a new subject they haven’t been trained on with over 90% accuracy using features as short as 12 seconds. Providing longer features enabled both models to reach an accuracy of 96%. This finding implies that not only the reflection coefficient carries information about the person’s stress, but that signals from different subjects share a common footprint of stress.

To further examine whether the signal sections recorded during “no stress” are inherently different from those recorded during the two stress periods, the same procedure was performed using the KMEANS2 unsupervised classifier. In this case, the model tried to separate the given features into two clusters without any ground truth labels and prior knowledge on the true nature of the features.

The unsupervised model was able to correctly separate the given test features into two distinct clusters. Unlike the supervised case, where the accuracy almost saturated after 36 seconds, the additional information improved the precision of the unsupervised model from 81% at 36 seconds to 90% at 60 seconds.

Next, the models were requested to classify the features to three classes – “no stress”, “mental stress”, and “physical stress”. The results of this multi-class classification are shown in Figure 2.

**Fig. 2:**
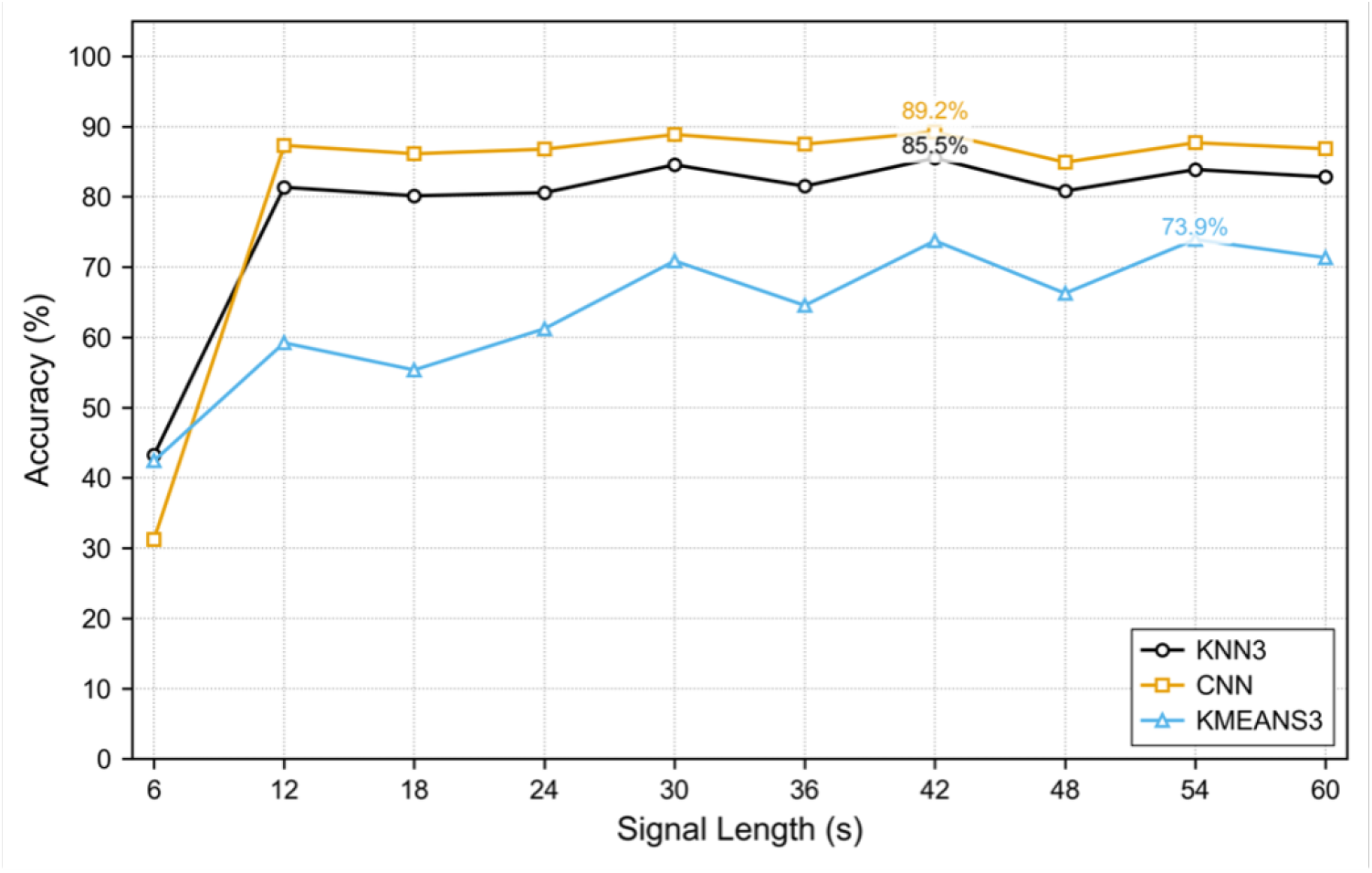
Multi-class classification accuracy of “no stress”, “mental stress” and “physical stress” features as a function of feature length using the supervised KNN3 and CNN classifiers, and the unsupervised KMEANS3 classifier.

Figure 2 shows that the supervised KNN3 and CNN models reached a maximal accuracy of 85% and 89%, respectively, with 42-second-long features. The unsupervised KMEANS3 model reaches 73% with 54-second-long features.

## 3 Discussion

Binary classification achieved an accuracy of over 90% using features as short as 12 seconds. The overall accuracy of the multiclass classification was lower than binary classification but reached over 80% accuracy with 12-second-long features and 89% with 42-second-long features.

Our findings show that the answer to both questions is positive, which signifies that the signal reflected from the skin in this frequency range carries inherent information about the person’s stress; information that is accessible via the EM signal itself without relying on any other physiological signals. Furthermore, this stress signature shares common characteristics between different subjects. Having this level of precision on the 12-second signals also indicates feasibility to use this approach for real-world applications of stress detection.

There are two potential reasons for the lower multiclass classification results. First, the physical stress section only lasted two minutes compared to the five minutes of mental stress. Therefore, to ensure a balanced training set, the number of features from each class had to be reduced to the lowest common denominator, which was the physical stress class. This resulted in fewer training features compared to binary classification.

In addition, when discussing the protocol with the participants after the experiment, it became apparent some fastened their grip on the ball but didn’t perform a repeated squeeze. This could have led to reduced physical stress compared to participants who did perform a repeated squeeze and to what was intended. Both hypotheses can be tested in future experiments by adjusting the experimental protocol.

These findings solidify the bind between the initial hypothesis, that the human sweat ducts function as biological antennas in the sub-THz range, and the subsequent hypothesis that the transmitted information should carry information about the person’s stress levels. As such, they form a steppingstone towards a remote and independent detection of stress and open a new non-invasive window into our nervous system.

## 4 Methods

### Participants

The participants were 21 healthy males and females, aged 20-30, recruited from the student body of The Hebrew University of Jerusalem. The experiment was approved by the Ethics Committee. All participants received an overview of the experiment and signed an informed consent form.

### Experimental Protocol

EM reflection coefficients were recorded from the palms of 21 participants for a period of 18 minutes. During the recording, the participants were subjected to a protocol designed to induce light mental and physical stress, via an interactive tablet-based application. The breakdown of the protocol is shown in Table 1.

**Table 1.** A breakdown of the experimental protocol with the corresponding class labels.

| Phase | Duration [min] | Binary Label | Multi. Label |
| --- | --- | --- | --- |
| Rest | 5 | 0 | 0 |
| Mental Stress | 5 | 1 | 1 |
| Rest | 5 | 0 | 0 |
| Physical Stress | 2 | 1 | 2 |
| Rest | 1 | 0 | 0 |

The participants were requested to avoid caffeine for two hours prior to the experiment. Upon arrival, they washed their hands with soap and were seated in front of the measurement table where a tablet computer with custom software was positioned in front of them at a comfortable height. For minimal distractions and interference, the protocol and the data collection were automated and after initiating the experiment, the experimenter left the room until the end of the protocol.

The experimental protocol consisted of several stress-free rest periods interleaved with mental and physical stress. Throughout the experiment, the software continuously instructed the participants on their actions. Accompanying music specifically tailored to each period was played using a set of headphones that the participants were given for a more immersive experience. The rest phase was repeated three times throughout the protocol: five minutes at the beginning of the protocol, to record the baseline of each participant, another five minutes after the mental stress stage for a reset period and one minute at the very end of the protocol. This stage consisted of watching a soothing video of a secluded tropical beach with the sound of the ocean waves in the headphones. It did not require any active participation.

The mental stress period was played immediately following the initial rest period. This stage lasted five minutes during which light mental stress was induced using color word tests and shape recognition games. This stage required active and quick participation. The time to provide an answer for each question was fixed and displayed as a count-down on the screen. It was also reduced continuously to prevent adaption and decrease in stress levels due to familiarity.

In the color word test, a name of a color appeared in the center of the screen written in colored letters with four answers written in black underneath it. The participant was requested to choose the answer that corresponded to the color of the letters of the word in the center (Figure 3B). This task is known to induce mental stress by creating an interference between the visual and semantic inputs in the brain, known as the Stroop effect [18, 19]

**Fig. 3:**
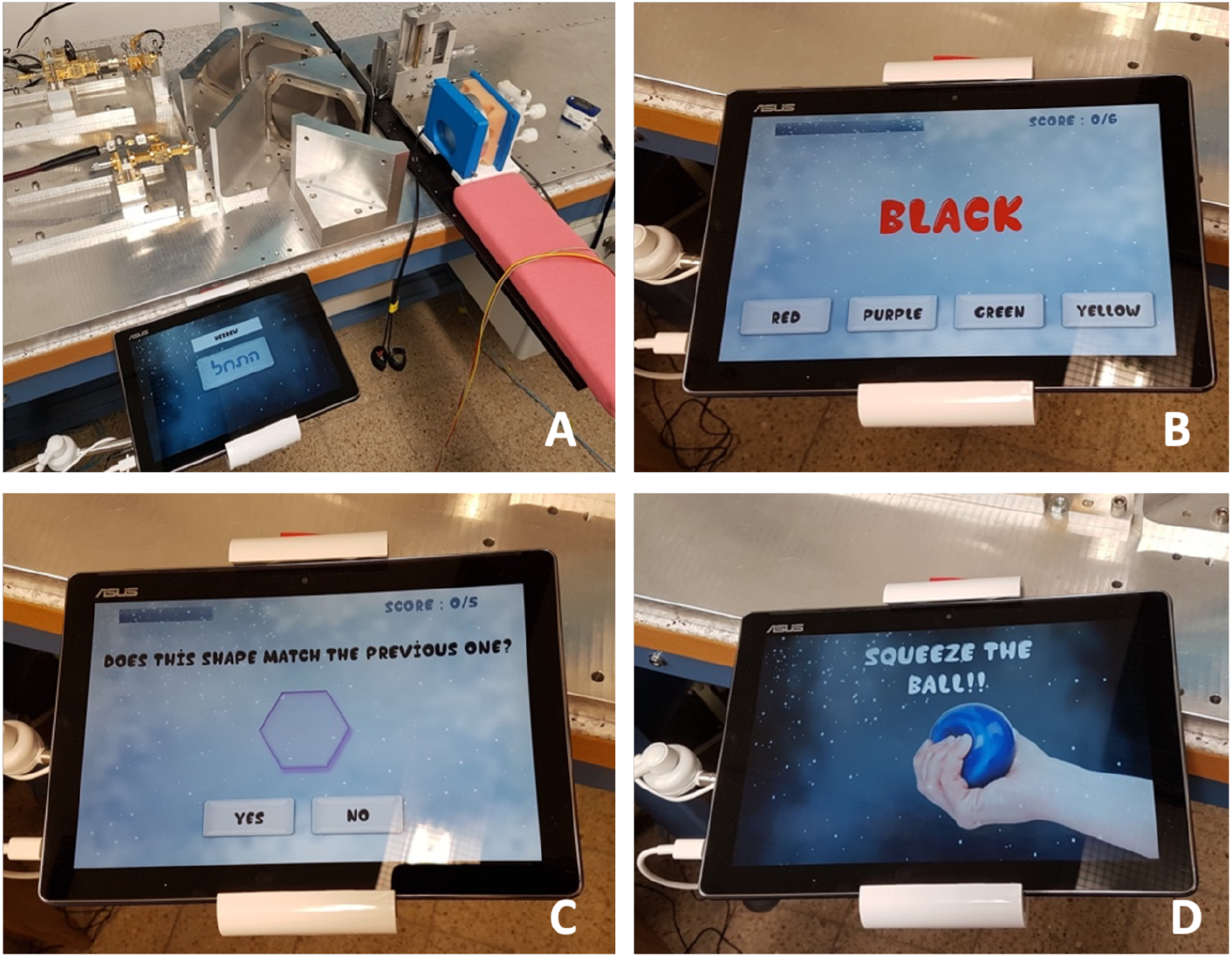
A. Top view of the measurement table with the tablet device and the hand rest. B. A sample screenshot of the color word test. C. A sample screenshot of the shape recognition test. D. A screenshot of the physical stress phase.

**Fig. 4:**
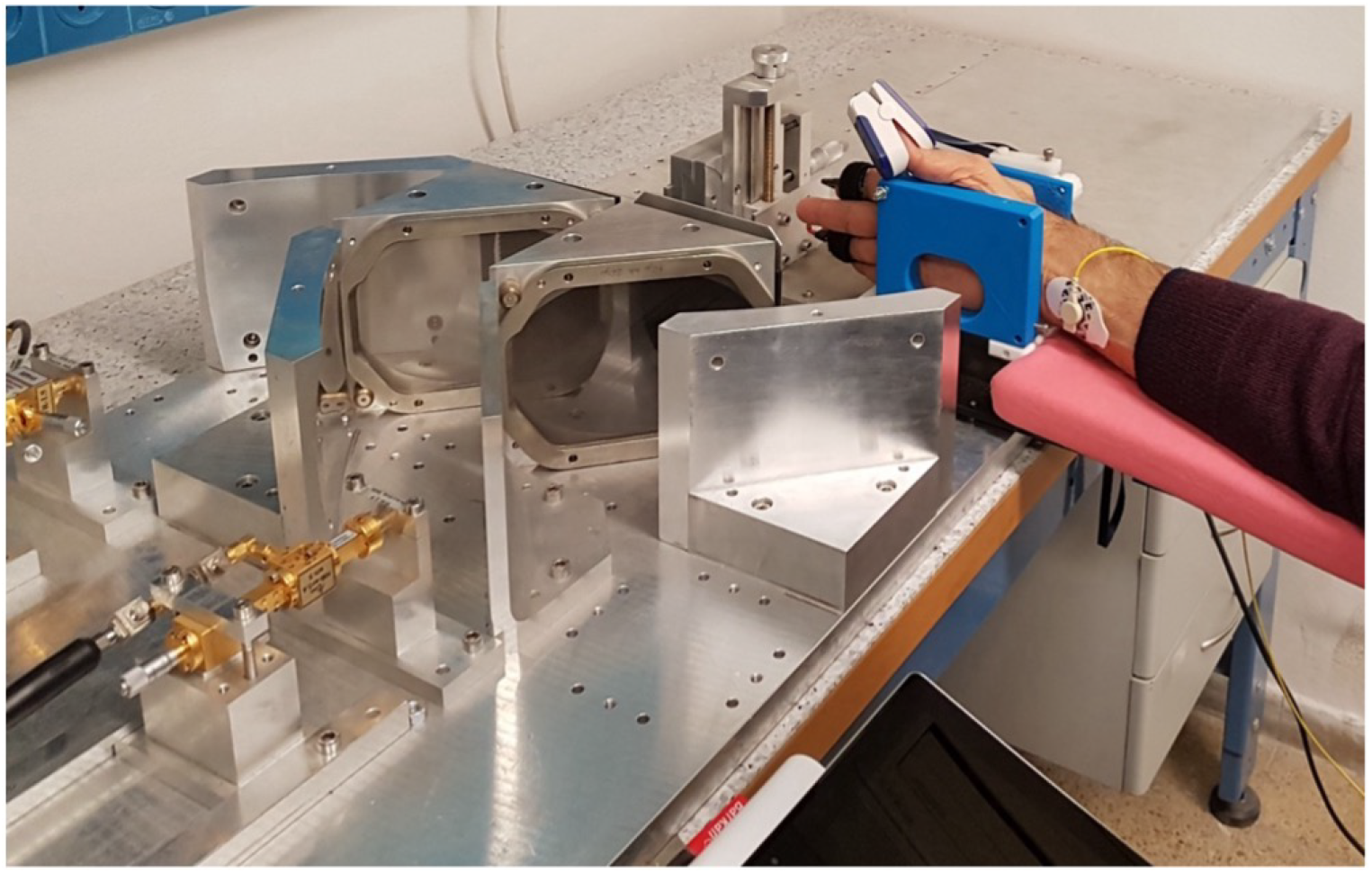
A demonstration of a hand fixed in the chassis on the measurement table.

During the shape recognition game, the participants were briefly shown a shape and then requested to say whether the shape on the following screen matched the initial one (Figure 3C). The shape on the second screen usually differed in color and size from the original shape.

The physical stress stage lasted two minutes, and during this stage the participants were instructed to repeatedly squeeze a rubber ball with their free left hand accompanied by upbeat workout music in their headphones. Apart from becoming increasingly more challenging as time progresses, this task is known to induce sweating due to physical activity in both the active palm as well as the palm that is not squeezing the ball, which is the hand that was recorded.

### EM Measurements

To measure the EM response from the skin throughout the experimental protocol, an EM signal in the frequency range of 426-432 GHz was generated by a Millimeter Vector Network Analyzer (MVNA-8-350, AB Millimetre, France). The signal was then focused on the right palm of the participant using a set of elliptical mirrors and grid polarizers, and the reflection coefficient from the palm was recorded by the detector of the MVNA unit. The full experimental setup is described in [10]. The right hand of the participant was positioned on the measurement table and held in place by a custom chassis at the focal point of the system. Due to the unique curvature of the palm, the hand position of each subject was further adjusted during the initial calibration stage to find the lateral and axial position with the strongest reflection.

Helical antennas can operate in the normal or axial mode where the generated radiation propagates perpendicularly or parallel to the helix, correspondingly. In the case of human sweat ducts, the mode that would generate a signal directed outwards from the palm is the axial mode. Previous simulation work [4] showed that the expected range for the axial mode lies between 360-520 GHz. Therefore, the central frequency selected for this experiment was 430 GHz. Strong attenuations and additional software limitations enforced a 6 GHz bandwidth around 429 GHz.

The MVNA was configured to perform a frequency sweep covering the selected range with 90 frequency points which were swept one after the other to obtain a continuous recording of the EM reflection from the skin throughout the experiment. A full sweep of all 90 frequencies took approximately 6 seconds.

In addition, to eliminate system artifacts, a calibration measurement was performed prior to every sequence of human measurements. This was done by measuring a flat metal plate, positioned in the chassis instead of the human hand, and recording the EM response for the full 18 minutes of the protocol.

### Data Analysis

The recorded data was processed to obtain train and test features for supervised and unsupervised ML models. Each raw EM recording was first divided by its corresponding metal recording to remove any system artifacts. In addition, as signals from different subjects can vary in amplitude, each recording was normalized to ensure the signals were comparable. The normalized signals were processed using Continuous Wavelet Transform (CWT) with a Mexican Hat to obtain a 2D representation that contains frequency information while preserving time information.

This full 18 minute, or 1080 second, long representation was sliced into multiple shorter features based on the tested duration (6-60 seconds). The slices examined were all multiples of 6 seconds, which is the duration of a full frequency sweep. This minimal time unit also allowed for a precise split of the stress periods to uniform sections (two minutes of physical stress can be split into exactly 20 features of 6 seconds or exactly 10 features of 12 seconds). There was no overlap between the distinct protocol phases within any given feature.

The 21 participants were randomly divided into train and test sets with a 70 to 30 percent ratio, and all the features of the train participants were fed into a classifier for the training procedure. In addition, as the physical and mental stress sections were shorter than the overall “no stress” period, to avoid bias in classification, the number of training features from each class was down sampled to the lowest common denominator: the number of physical stress features in the multi-class classification and the overall number of stress features in the binary classification.

Two classical supervised models as well as one unsupervised model were examined. The chosen classical supervised classifiers were K Nearest Neighbors with three neighbors (KNN3) and a Convolutional Neural Network (CNN). The unsupervised model was KMEANS with two (KMEANS2) or three clusters (KMEANS3).

The models were trained and tested on binary classification between “stress” and “no stress” (rest) as well as multiclass classification between “mental stress”, “physical stress”, and “no stress”. The classification results for each of the classifiers were averaged over 50 independent executions. Each execution randomized a different set of test subjects, but the same randomization seeds were used across all classifiers to ensure comparable results. K-fold validation was used during the training process to ensure that the model did not overfit any particular subset of subjects.

## 5 Author Contributions

AKG, YG, YF, SE and PBI conceived the study. AKG developed the protocol and performed the measurements. AKG and YG processed the data. AKG and YG wrote the manuscript. YG and AKG generated the figures. SE handled ethics approval. All authors reviewed and commented on the manuscript.

